# Diversified Diet Feeding Practice is Low Compared to the WHO Recommendation in the Dabat Demographic and Health Surveillance System Site: Finding from the Baseline Survey of Nutrition Project, 2016

**DOI:** 10.1101/553875

**Authors:** Zegeye Abebe, Amare Tariku, Gashaw Andargie Bikes, Molla Mesele Wassie, Kedir Abdella, Tadesse Awoke, Ejigu Gebeye, Azeb Atnafu Gete, Melkie Edris Yesuf, Yigzaw Kebede, Kassahun Alemu, Abebaw Addis, Esmael Ali Muhammad, Solomon Mekonnen Abebe, Aysheshim Kasahun belew, Melkamu Tamir, Melkitu Fentie, Adane Kebede, Kindie Fentahun

## Abstract

**Introduction:** Improving infant and young child feeding practices is critical to improved nutrition, health, and development of children. The country of Ethiopia has also adopted the WHO recommendations of child feeding practices and developed the national guideline of infant and young child feeding to improve child’s nutrition and health status. However, a few children start and received appropriate complementary feeding based on the recommendation. Therefore, the study aimed to determine dietary diversity score and its associated factors among under five children at Dabat Demography Surveillance System site (HDSS), northwest Ethiopia.

**Methods:** A cross-sectional community based study was from February to June 2016. All children aged 6-59 months old who lived in HDSS site were included in the survey. The collected data were checked and entered into Epi info version 7 and exported to STATA version 14 statistical software for analysis. Both crude odds ratio (COR) and adjusted odds ratio (AOR) with the corresponding 95% confidence interval (CI) was calculated to show the strength of association. Finally, a p-value of 0.05 was used to determine if the association was statistically significant.

**Results:** In this study, about 34.87% (95%CI: 33.27, 36.49%) of the children received adequately diversified diet. The odds of receiving adequately diversified diet was higher among children whose mother had secondary and above education (AOR= 6.51; 95%CI: 4.95, 8.56), mother who had ANC (AOR = 1.90; 95%CI: 1.60, 2.26) and PNC visit (AOR= 1.31; 95%CI: 1.00, 1, 72). However, a lower dietary diversity score is observed among young children (AOR=0.59; 95%CI: 0.41, 0.85), and children from food inscured household (AOR=0.76; 95%CI: 0.63, 0.92).

**Conclusions:** Diversified diet feeding practice is low compared to the WHO recommendation in the surveillance site. Age of the child, maternal education, ANC and PNC visit, and household food insecurity were significantly associated with Dietary diversity score of children. Hence, various actions need to scale up the current practices of child feeding by improving HHFSS, strengthening ANC and PNC counselling about child feeding options, and feeding of young infants.

## Introduction

Malnutrition, in its various form, remains a pressing and significant health problem of children in Ethiopia (1). It affects mortality and ill-health along the entire continuum of care from early childhood to adulthood (2). It is thus clear that the prevention of young child undernutrition is a long-term investment that will benefit the current generation as well as their children (3). The first two years of life is called a critical period to ensure the child’s development through optimum feeding practices (4, 5). If children are undernourished before they reach the age of 2 years, they could suffer irreversible physical and mental damage and this will undoubtedly influence their future health and wellness. Thus, improving infant and young child feeding practices is therefore critical to improved nutrition, health, and development of children (2, 6).

It is confirmed that exclusive breast-feeding adequately provides for children’s energy and nutrient needs in the first six months of life. However breast-milk alone cannot meet the increased energy and nutrient requirements as children get older (7). Thus, the World Health Organization (WHO) recommends that infants should be exclusively breastfed for the first 6 months of life, after which they are introduced to appropriate complementary foods as they continue to breastfeed (6). Complementary food introduction should be timely, nutritionally adequate, appropriate, and safe for the development of children’s full human potential. In addition, it should include a variety of foods; grains, roots and tubers; Legumes and nuts; dairy products; flesh foods (meats/fish/poultry); eggs; vitamin A-rich fruits and vegetables; and other fruits and vegetables to ensure daily nutritional requirements of children (8).

Despite dietary diversity is a potentially and useful indicator for nutrient adequacy of a diet consumed by an individual and their nutritional status (9–11), complementary feeding continues as a challenge to good nutrition in children. For instance, in Nigeria 26.5% of under five children received adequately diversified diet (12), in Kenya 76% attained minimum meal frequency; 41% of the children attained a minimum dietary diversity; and 27% attained minimum acceptable diet (13). Similarly, inappropriate complementary feeding is a common practices in Ethiopia. According to the EDHS 2016 report, 7% of the children received the minimum feeding standards, 14% received adequate DDS and 45% the recommended meal frequency. Similarly, previously studies in Ethiopia also noted that feeding of diversified diet is well under the recommendation, 25% in Gondar, 35% in Bihar Dar, and 45% in Addis Ababa. In addition, the country has high burden of child undernutrition, 38% children were stunted and 10 were wasted (1).

To improve child feeding practices and nutritional status of children, the government of Ethiopia has been conducted different activities. Accordingly, the country adopted the WHO recommendations of child feeding practices and developed the national guideline of IYCF to improve child’s nutrition and health status (14). However, a few children start and received appropriate complementary feeding based on the recommendation. Therefore, investigating the determinate of IYCF practices are important to continuously monitor and provide evidence-based decision-making in interventions designed to improve child feeding practices and reduce undernutrition. Also, there is need to update the existing literatures using recent and representative surveillance data.

## Methods

### Study setting and area

A cross-sectional community based study was conducted in one of the surveillance site of the University of Gondar from February to June 2016. The HDSS is found in Dabat district, northwest Ethiopia. It is located at 814 km away from Addis Ababa, the capital city of Ethiopia. The district has 6 health centers, 31 rural and 5 urban health posts with a total populations of 185,307. The HDSS site has been running since 1996, and hosted by the University of Gondar. The surveillance site covers thirteen randomly selected kebeles (four urban and nine rural kebeles) in different ecological zones (high land, middle land, and low land).

### Study participants, Data collection tool and procedures

All children aged 6-59 months old who lived in HDSS site were included in the survey. The data were collected from mothers of children using a structured questionnaire via interview. Questioners were adopted from previous studies with some modification to fit the local context. The questionnaire was first developed in English and translated into Amharic then back translated into English to maintain consistency. Pretest was done on five percent participants out of the study area. The data were collected by the HDSS site data collectors. A two days training was given for ten and three data collectors and supervisors, respectively. During data collection intensive supervision was carried out by investigators and supervisors.

### Variable measurements

#### Timely introduction of solid, semi-solid or soft foods

Mothers were asked to indicate if the target child had initiated complementary food within 6-8 months of age. Then, Infants 6–8 months of age who received solid, semi-solid or soft foods within the stated time is considered as the appropriate time for his/her age.

#### Minimum dietary diversity

The children’s complementary food dietary diversity score (DDS) definition was based on seven food groups, including grains/roots/tubers; legumes and nuts; dairy products; flesh foods (meats/fish/poultry); eggs; vitamin A-rich fruits and vegetables (VAFV); and other fruits and vegetables (OFV). The DDS was constructed by assigning one point to each of the defined food groups, for a maximum of seven points. Then, we defined minimum complementary food dietary diversity as consumption of food from at least four different food groups (DDS≥4).

#### Household food security status

This was assessed by using the standardized questionnaire developed by Food and Nutritional Technical Assistance (FANTA). A one month recall period with two types of questions, nine occurrence questions followed by three frequency of occurrence questions for each event, was used to determine the status. Finally, households were categorized as food secured when the score was ≤1, and food insecure for a score ≥2.

### Data processing and analysis

The collected data were checked and entered into Epi info version 7 and exported to STATA version 14 statistical software for analysis. Descriptive statistics were carried out and the result was presented using text, and tables. A binary logistic regression model was fitted to identify factors associated with dietary diversity score. Variables with a p-value of less than 0.2 in the bivariable analysis were fitted into the multivariable logistic regression analysis. Both crude odds ratio (COR) and adjusted odds ratio (AOR) with the corresponding 95% confidence interval (CI) was calculated to show the strength of association. Finally, a p-value of 0.05 was used to determine if the association was statistically significant.

### Ethical approval

Ethical clearance was obtained from the Institutional review board of the University of Gondar. Informed verbal consent was obtained from each participant.

## Results

### Characteristics of the parents and their children

In this study, a total of 3433 children were included. Almost half, (49.2%), of the children were male and one fourth of the children were in the age range of 36-47 months. Regarding maternal characteristics, 70.3% were unable to read and write followed by primary education (19.84%), 86.13% were housewife, and 85.49% were married (Table 1).

**Table 1:** Characteristics of the parents and their children at Dabat HDSS site, northwest Ethiopia, May to August 2016 (n= 3433)

| Variables | Frequency | Percentage |
| --- | --- | --- |
| **Sex of the child** |  |  |
| Male | 1689 | 49.20 |
| Female | 1744 | 50.80 |
| **Child age** |  |  |
| 6-11 | 371 | 10.81 |
| 12-23 | 802 | 23.36 |
| 24-35 | 774 | 22.55 |
| 36-47 | 839 | 24.44 |
| 48-59 | 647 | 18.85 |
| **ANC visit** |  |  |
| No | 1202 | 35.01 |
| Yes | 2231 | 64.99 |
| **Maternal education** |  |  |
| Unable to read and write | 2404 | 70.03 |
| Primary education | 681 | 19.84 |
| Secondary and above | 348 | 10.14 |
| **Mother's occupation** |  |  |
| Farmer | 624 | 18.18 |
| Housewife | 2333 | 86.13 |
| Merchant | 286 | 8.33 |
| Others | 190 | 5.53 |
| **Maternal status** |  |  |
| Single | 323 | 9.41 |
| Married | 2935 | 85.49 |
| Other | 175 | 5.10 |
| **Residence** |  |  |
| Urban | 588 | 17.13 |
| Rural | 2845 | 82.87 |
| PNC visit |  |  |
| No | 3153 | 91.84 |
| Yes | 280 | 8.16 |

### Feeding practices and Family food choice to their children

In this study, about 34.87% (95%CI: 33.27, 36.49%) of the children received adequately diversified diet. Of all children, 5.62% started complementary feeding early before sex months and 59.63% started complementary feeding at six months. About 18.64% of the children received meat and flesh foods 24hr before the date of survey. Similarly, consumption of foods rich in vitamin A or iron remains low among children in the study area. About 27.06% of children consumed foods rich in vitamin A, and 18.64% consumed iron rich foods 24 hours before the interview. Around 60% of the children started complementary feeing at the age of six months (Table 2).

**Table 2:** Feeding practices and family food choice for their children at Dabat HDSS site, northwest Ethiopia, 2017 (n=3433)

| Variables | Frequency | Percentage |
| --- | --- | --- |
| **Breast feeding status** |  |  |
| No | 1,640 | 47.77 |
| Yes | 1,793 | 52.23 |
| **Time of complementary introduction** |  |  |
| Before six months | 193 | 5.62 |
| At six months | 2047 | 59.63 |
| After six months | 1193 | 34.76 |
| **Preparation of CF** |  |  |
| For the child only | 832 | 24.24 |
| With the family | 2601 | 75.76 |
| **DDS** |  |  |
| Adequate | 1197 | 34.87 |
| Inadequate | 2236 | 65.13 |
| **Child feeding style** |  |  |
| With the family | 1395 | 40.64 |
| Separately | 2038 | 59.36 |
| **Food source** |  |  |
| Home garden | 2483 | 73.83 |
| Market | 949 | 27.95 |
| **Frequency of food buying (n=1180)** |  |  |
| Daily | 20 | 1.69 |
| 2-3 times weekly | 95 | 8.05 |
| Weekly | 269 | 22.80 |
| Once in two weeks | 180 | 15.25 |
| Monthly | 616 | 52.21 |
| We didn't buy | 20 | 1.69 |
| **Reasons not to buy food (n=11)** |  |  |
| Too costly | 4 | 36.36 |
| The money sent for other purpose | 3 | 27.27 |
| The food is unavailable | 2 | 18.18 |
| Others | 2 | 18.18 |
| **Distance b/n home and the market (n=1183)** |  |  |
| <1km | 589 | 49.79 |
| 1-4km | 216 | 18.26 |
| 5-10km | 27 | 2.28 |
| 11-20km | 95 | 8.03 |
| 21-30km | 89 | 7.52 |
| 31-40km | 114 | 9.64 |
| 41-50km | 30 | 2.54 |
| >50km | 23 | 1.96 |
| **Transportation to market** |  |  |
| On foot | 1173 | 99.66 |
| Others (car and animals) | 4 | 0.34 |
| **Food preference/ Food choice to their children** |  |  |
| No | 2794 | 81.40 |
| Yes | 639 | 18.60 |
| **Types of Food preference to their children (n=639)** |  |  |
| Fruit and vegetables | 325 | 50.8 |
| Meat and fish | 447 | 69.95 |
| Egg, milk and milk product | 476 | 74.49 |
| Honey | 262 | 41.00 |
| Packed foods | 306 | 47.89 |
| Cereals and food made from it | 582 | 91.08 |

### Factors associated with DDS among children

Table 3 shows factors associated with appropriate complementary feeding practices of children aged 6–59 months, a bivariate and multivariate analyses. Accordingly, age of the children, maternal education, maternal ANC and PNC follow up, household food security status and method of child feeding were associated with DDS.

**Table 3:** Factors associated with DDS among children at Dabat DHSS site, northwest Ethiopia, 2017

| Variables | DDS |  | COR 95%CI | AOR 95%CI |
| --- | --- | --- | --- | --- |
|  | Adequate | Inadequate |  |  |
| **Age of children** |  |  |  |  |
| 6-11 | 113 | 258 | 0.73 (0.56, 0.96) | **0.59 (0.41, 0.85)** |
| 12-23 | 270 | 532 | 0.85 (0.68, 1.05) | 0.78 (0.58, 1.06) |
| 24-35 | 270 | 504 | 0.89 (0.72, 1.11) | 0.80 (0.61, 1.04) |
| 36-47 | 302 | 537 | 0.94 (0.76, 1.16) | 0.90 (0.71, 1.13) |
| 48-59 | 242 | 405 | 1.00 | 1.00 |
| **Sex of the children** |  |  |  |  |
| Male | 610 | 1079 | 1.00 | 1.00 |
| Female | 587 | 1157 | 0.89 (0.78, 1.03) | 0.92 (0.79, 1.07) |
| **Mother's education** |  |  |  |  |
| Unable to read and write | 635 | 1769 | 1.00 | 1.00 |
| Primary education | 304 | 377 | 2.25 (1.88, 2.68) | **2.11 (1.76, 2.53)** |
| Secondary and above | 258 | 90 | 7.98 (6.18, 10.32) | **6.51 (4.95, 8.56)** |
| **ANC visit** |  |  |  |  |
| No | 278 | 924 | 1.00 | **1.00** |
| Yes | 919 | 1312 | 2.33 (1.99, 2.73) | **1.90 (1.60, 2.26)** |
| **PNC visit** |  |  |  |  |
| No | 1067 | 2086 | 1.00 | 1.00 |
| Yes | 130 | 150 | 1.69 (1.32, 2.16) | **1.31 (1.00, 1.72)** |
| **Currently breastfeeding** |  |  |  |  |
| No | 606 | 1034 | 1.19 (1.04, 1.37) | 1.04 (0.84, 1.29) |
| Yes | 591 | 1202 | 1.00 | 1.00 |
| **Methods of child feeding** |  |  |  |  |
| With family | 526 | 869 | 1.23 (1.07, 1.42) | **1.39 (1.17, 1.65)** |
| The child alone | 671 | 1367 | 1.00 | **1.00** |
| **Methods of CF preparation** |  |  |  |  |
| Separately | 323 | 509 | 1.00 | 1.00 |
| With the family | 874 | 1727 | 0.80 (0.69, 0.94) | 1.00 (0.81, 1.23) |
| **Food security** |  |  |  |  |
| Secured | 994 | 1714 | 1.00 | **1.00** |
| Unsecured | 203 | 522 | 0.67 (0.56, 0.80) | **0.76 (0.63, 0.92)** |
| **Food preference** |  |  |  |  |
| No | 954 | 1840 | 1.00 | 1.00 |
| Yes | 243 | 396 | 1.18 (0.99, 1.41) | 1.12 (0.92, 1.36) |

The odds of receiving adequately diversified diet among younger children (6–11 months) was 41% (AOR=0.59; 95%CI: 0.41, 0.85) less likely compared with older children (48-59 months). Mothers who had secondary and above educational status were 6 times (AOR= 6.51; 95%CI: 4.95, 8.56) more likely to provide adequately diversified diet to their children compared to mothers who had no education. Similarly, mothers who attended primary school were about 2 times (AOR= 2.11; 95%CI: 1.76, 2.53) more likely to practice adequate diversified diet than those who had no formal education. Mothers who had ANC and PNC follow up were about 2 times more likely to provide diversified diet than those who had no ANC and PNC follow up, AOR = 1.90; 95%CI: 1.60, 2.26 and AOR= 1.31; 95%CI: 1.00, 1,72, respectively.

Finally, those children from food insecured household were 24% less likely to received adequately diversified diet compared to children from food secured household (AOR=0.76; 95%CI: 0.63, 0.92).

## Discussion

Proper feeding of infants and young children is necessary to ensure normal growth and development. Diversified diet, which provides all the essential nutrients in sufficient quantities to meet the children needs, is an adequate or balanced diet (15).

The current study found that only 34.87% of the children received diversified food based on the recommendation. This indicates various and rigorous actions needed to scale up for appropriate child feeding practices. Because complementary foods are expected to have all important nutrients and should be timely, adequate, safe and appropriate to tackle the most pressing nutritional problem of the children (16). In addition, adequate intake of micronutrient, particularly, iron and vitamin A rich foods, has an impact on physical and mental development, strong immune function and blood cell formation. However, this finding is higher than a report from Southern region, 18.8% (8) and Benishangul Gumuz Region, 23.7% (17) of Ethiopia. This might be explained by the disparity in educational, maternal health services utilization, particularly ANC and PNC, socioeconomic and cultural differences between the study settings.

Mothers who had ANC and PNC follow up were more likely to provide diversified diet to their children. This finding is supported with other study in Ethiopia (18, 19). ANC and PNC follow up may supported with nutrition education, which is a very important input to help people to select an adequate and variety of diet. Mothers can learn how to select an adequate, so simple, and attractive diet to their children. In addition, nutrition education may encouraged mothers to retain the existing beneficial food habits and add other foods which may help to meet their children nutritional needs, healthy feeding practices, offering of energy dense and adequate amount of complementary foods (20). Successful nutrition education reinforces the existing cultural pattern and brings about qualitative improvement by using available food resources (21). A maternal education level, primary education and above was significantly associated with dietary diversity score of children; educated mothers more likely to provide adequately dietary diversified diet to their children compared to those mothers who were unable to read and write. It is clear that education is the primary intervention in all aspects of health promotion and prevention of health related problems. Specifically, maternal education is the corner stone of proper child growth and development, appropriate child feeding and prevention of hygiene related diseases (2). In addition, educated mothers can easily understand the nutrient requirement of their children and they can also plan, select and prepared energy dense and adequate complementary foods to their children.

The age of the child is also found to have an association with dietary diversity score. Younger children were less likely to receive the recommended diversified diet compared to the older one. This finding is supported by that of another study conducted in Ethiopia (19, 22). This is probably because younger infants’ are mostly breastfed, so the need for a frequent feeding of extra solid food is not perceived as important or a priority by mothers and caretakers for feeding infants of this age. In addition, older children have the chance of eating family diet as it is observed from this study.

The likely hood of having adequate dietary diversity was higher among children who feed with their family compared to children feed alone. This is because family characteristics such as parental role modeling for eating, parental encouragement, parents’ food preferences, regular meal patterns and feeding practices encourage a higher intake of diversified diet among children (23, 24).

Finally, household food insecurity is associated with dietary diversity score of children. This finding is supported with other studies (13, 25). It is known that HHFSS is one of the underlined cause of child malnutrition and hunger. Food inscured households may affect time of introduction of complementary feeding and adherence to the infants and young child feeding recommendation. Therefore, low dietary diversity may be due to limited income available to purchase foods, and reducing the variety of foods consumed and preparing cereal based monotones diet are the coping strategies adopted in the face of food insecurity (26).

In conclusion, diversified diet feeding practice is low compared to the WHO recommendation. Hence, various actions need to scale up the current practices of child feeding by improving HHFSS, strengthening ANC and PNC counselling about child feeding options, and feeding of young infants.

## Abbreviations

HDSS: Health and Demographic Surveillance System
EDHS: Ethiopia Demography and Health Survey
WHO: World Health Organization
BMI: body mass index
FANTA: Food and Nutrition Technical Assistance
AOR: adjusted odd ratio
CI: confidence interval

## Authors’ contributions

Conceptualization: AT, KAG, GAB, KA, AKB, MMW, TA, EG, ZA, AAG, MEY, YK, AAG, EAM, SAM, MF, AK and KF.

Data curator: AT, KAG, GAB, KA, AKB, MMW, TA, EG, ZA, AAG, MEY, YK, AAG, EAM, SAM, MF, AK and KF.

Formal analysis: AT, KAG, GAB, KA, AKB, MMW, TA, EG, ZA, AAG, MEY, YK, AAG, and KF.

Funding acquisition: AT, KAG, GAB, KA, AKB, MMW, TA, EG, ZA, AAG, MEY, YK, AAG, EAM, SAM, MF, AK and KF.

Investigation: AT, KAG, GAB, KA, AKB, MMW, TA, EG, ZA, AAG, MEY, YK, AAG, EAM, SAM, MF, AK and KF.

Methodology: AT, KAG, GAB, KA, AKB, MMW, TA, EG, ZA, AAG, MEY, YK, AAG, EAM, SAM, MF, AK and KF.

Resources: AT, KAG, GAB, KA, AKB, MMW, TA, EG, ZA, AAG, MEY, YK, AAG, EAM, SAM, MF, AK and KF.

Software: AT, KAG, GAB, KA, AKB, MMW, TA, EG, ZA, AAG, MEY, YK, AAG, and KF.

Supervision: AT, KAG, GAB, KA, AKB, MMW, TA, EG, ZA, AAG, MEY, YK, AAG, and KF.

Validation: AT, KAG, GAB, KA, AKB, MMW, TA, EG, ZA, AAG, MEY, YK, AAG, EAM, SAM, MF, AK and KF.

Visualization: AT, KAG, GAB, KA, AKB, MMW, TA, EG, ZA, AAG, MEY, YK, AAG, EAM, SAM, MF, AK and KF.

Writing—original draft: AT, KAG, GAB, KA, AKB, MMW, TA, EG, ZA, AAG, MEY, YK, AAG, and KF.

Writing—review and editing: AT, KAG, GAB, KA, AKB, MMW, TA, EG, ZA, AAG, MEY, YK, AAG, and KF.

## Acknowledgements

We are thankful to adolescent girls, interviewed families, data collectors and supervisors of this study.

## Availability of data and materials

Data will be available upon request from the corresponding author.

## Consent for publication

Not applicable.

